# Adult crowding induces sexual dimorphism in chronic stress-response in *Drosophila melanogaster*

**DOI:** 10.1101/702357

**Authors:** Shraddha Lall, Akhila Mudunuri, S. Santhosh, Akshay Malwade, Aarcha Thadi, Gayathri Kondakath, Sutirth Dey

## Abstract

Stress-induced mood disorders such as depression and anxiety are sexually dimorphic in human beings. Studying behavioural stress-responses in non-human animal models can help better understand the behavioural manifestations of these disorders and the dimorphism in their prevalence. Here we explore how sexes show differential behavioural responses to different chronic stressors, both abiotic and biotic, by using outbred populations of *Drosophila melanogaster*. The behaviours studied – namely, anhedonia, motivation to explore a novel habitat, locomotor activity and sleep levels – have been well-investigated in human and rodent-based models of stress disorders. These behaviours were studied in the context of two different stressors – mechanical perturbation and adult crowding. Responses to stress were found to be sexually dimorphic, and stressed females showed more behavioural changes, such as a reduced motivation to explore a novel habitat. Furthermore, adult crowding caused a greater number of sexually dimorphic behavioural changes than mechanical perturbation. For instance, while mechanical perturbation caused anhedonia across sexes, only females were anhedonic after crowding. We thus make a case for *Drosophila melanogaster* as a model system for studying sexual dimorphism in stress-induced mood disorders in humans.

**SUMMARY STATEMENT:** Female fruit flies, like their human counterparts, are more prone to chronic stress-induced mood disorders like anhedonia or reduced activity. This sexual dimorphism was more evident in a biotic stress.

## INTRODUCTION

Stress-induced mood disorders (SIMDs) such as depression (Krishnan and Nestler, 2008; Van Praag, 2004) and anxiety (Shin and Liberzon, 2009) can cause debilitating psychological symptoms including suicidal tendencies, loss of sleep or appetite and reduced interest in pleasurable activities (World Health Organisation, 2017; see Cryan and Holmes, 2005; Wong and Licinio, 2001 for reviews). Moreover, they have been etiologically associated with other physiological ailments like type-2 diabetes (Knol et al., 2006), cardiac disease and cerebrovascular disease (reviewed in Evans et al., 2005). In order to better manage and treat these disorders, identifying therapeutic targets and drugs for SIMDs or enhancing the efficacy of the current treatments (Wong and Licinio, 2004) has been a key focus for researchers for over 60 years. Much of the research in this area has used animal models to investigate the underlying symptoms and the predisposition to these disorders, as well as develop novel therapeutic strategies. Mammals are often seen as natural models in which to study the stress-response, with rodent (see Abelaira et al., 2013; Cryan and Holmes, 2005; Willner, 2017; Willner et al., 1992 for reviews), dog (Seligman and Maier, 1967) and primate (see Mendoza et al., 2000 for a discussion) models being popular. Recently, it has been shown that invertebrates, specifically *Drosophila melanogaster*, can also be a useful model-system for studying SIMDs (reviewed in Iliadi, 2009). Unfortunately, certain well-known features of human SIMDs remain relatively less explored in the model systems.

One of the features of SIMDs in humans is that they can be sexually dimorphic, i.e. males and females can differ in terms of how they are affected by these disorders. For example, in humans, the prevalence of depression (Kessler, 2003; Nolen-Hoeksema, 1987) and generalized anxiety disorder (McLean et al., 2011; Wittchen et al., 1994) is reported to be twice as much in females than in males. Furthermore, it is known that males and females can respond differently to drugs used for treating SIMDs (reviewed in Frackiewicz et al., 2000). Curiously though, in spite of this, how sex affects SIMDs has remained relatively less investigated in both vertebrate (see Palanza and Parmigiani, 2017 for a discussion) and invertebrate model systems (however, see Neckameyer and Matsuo, 2008; Neckameyer and Nieto-Romero, 2015)

To complicate matters further, the method of stress-induction in animal models can affect the ensuing behavioural changes. For instance, while acute oxidative stress for 24 hours caused a decrease in exploratory locomotion in fruit flies across most ages and sexes, starvation both increased and decreased exploration depending upon the age and sex of the fly (Neckameyer and Matsuo, 2008). However, when a combination of stressors like starvation, heat stress, cold stress and sleep deprivation was used over ten days, no change was observed in exploratory behaviour in male flies (Araujo et al., 2018). The same chronic stress protocol led to no change in short-term locomotor behaviour over 1-2 minutes (Araujo et al., 2018). However, when a 3-day long chronic vibrational stress protocol was used, reduced activity over a 15-minute period was observed in male flies (Ries et al., 2017). Further, when a paradigm of learned helplessness was used, wherein flies were given electric shocks in a 10-20-minute period, they showed an increased latency to escape from the shock-box after the stress (Batsching et al., 2016). However, there were no long-term changes in locomotor behaviour in an open-field arena, thus making this stressor environment specific (Batsching et al., 2016).

Apart from the abiotic components mentioned above, the social environment of a species can also act as a potential source of stress (Palanza, 2001). For example, social isolation for 24-hours in fruit flies has been shown to reduce the number of transitions in a light-dark box, regardless of the age and sex of the fly, thus suggesting a negative impact of this acute stressor on exploratory behaviour (Neckameyer and Nieto-Romero, 2015). Adult crowding for 3 days has been shown to reduce both mortality during crowding and post-stress fecundity in fruit flies (Joshi et al., 1998). Furthermore, adult crowding in flies is known to lead to a reduction in lifespan, possibly due a reduction in stored energy reserves (Joshi and Mueller, 1997).

In this study, we attempt to understand how sexual dimorphism in stress-response is modulated by the nature of the stressor in the common fruit fly *Drosophila melanogaster*. For this purpose, we studied an abiotic stressor (namely, mechanical perturbation) as well as a biotic one (namely, adult crowding). We employed 3-day long chronic stress protocols, and provided overnight rest before any behavioural measurements. We measured the stress-response in terms of three different behaviours – anhedonia, exploratory behaviour, and locomotor activity/sleep. Anhedonia – a reduction in normally rewarding, pleasurable activities – is a core symptom of depression in human beings (Krishnan and Nestler, 2008; Nestler et al., 2002), and has been previously observed in stressed male fruit flies (Araujo et al., 2018; Ries et al., 2017). Similarly, the tendency to explore a novel arena is influenced by the motivational state of the fly (Liu et al., 2007), and edge-preference and reduced exploration of an arena is seen as a marker of shelter-seeking (Liu et al., 2007). Such a reduction in motivation to explore and investigate a novel arena has also been observed in stressed rodents (Strekalova et al., 2004; Willner, 1997). Finally, altered psychomotor activity (Nelson and Charney, 1981), insomnia (or a lack of sleep) and hypersomnia (excessive sleeping) have been diagnostic features for SIMDs in humans (Nutt et al., 2008). To check whether stress causes any changes in psychomotor activity and rest levels, we recorded the locomotor behaviour of the flies over a 6-hour period. We measured both the amount of time the fly spends resting or sleeping, as well as the activity level of the fly during wakefulness.

We found that male and female fruit flies responded differently to the two stressors, with adult crowding leading to a larger number of sexually dimorphic behavioural changes. Both males and females, after either stress, showed increased levels of sleep. However, while mechanical perturbation caused anhedonia and made flies hyperactive across sexes, the changes in these behaviours was sexually dimorphic after adult crowding. Further, female flies, across stressors, showed a reduced motivation to explore a novel arena, while male flies did not. Thus, we conclude that sex plays a crucial role in modulating the behavioural stress-response in fruit flies. We finally discuss the impact of these results on modelling stress-responses in light of the existent sexual dimorphism in SIMDs in humans.

## MATERIALS AND METHODS

### Experimental Populations

For the set of experiments on ancestral non-selected flies, a laboratory-bred baseline population of *Drosophila melanogaster* (DB_4_) was used (breeding population of ~2400, 21-day discrete generation cycle). The detailed maintenance protocol of this population can be found elsewhere (Sah et al., 2013). For each assay, age-matched flies were used for all treatment groups. Adult flies, between 11 and 13 days old, were separated by sex under light CO_2_ anaesthesia. They were subjected to the experimental protocol after allowing them to recover overnight.

### Stressors

#### Mechanical perturbation

This stress paradigm was modified from the vibrational stress protocol in Ries et al., (2017). 25-50 flies of either sex were kept in vials containing a sponge at the bottom, soaked with water, for the duration of the stress. The treatment vials were placed on a platform shaker, rotating at 400 RPM, while the control vials were placed on an undisturbed surface. The mechanical perturbation was provided for 15 minutes, followed by a period of rest for 15 minutes. This was repeated over the entire duration of the stress protocol, which was 8 hours for males and 10 hours for females. These durations were finalized based on standardizations for both sexes, to ensure that there was minimal mortality due to the stress (< 3%). They were then transferred to vials containing food and allowed to recover overnight. The same protocol was carried out at the same time of the day for 3 days; on the 4^th^ day, the flies were subjected to various assays.

#### Adult crowding

The protocol was modified from Joshi and Mueller, (1997). 150 flies of either sex were placed in a vial with ~6mL of food. A sponge plug was pushed into the vial such that there was 0.7cm distance between the food and the plug for males and 1cm for females. This stress was maintained for 72 hours at a stretch. After this, the flies were transferred to round-bottom fly bottles with food and the crowding stress was relaxed for 14 hours before the assays were conducted. Control vials had 50 flies of either sex, maintained under uncrowded conditions (i.e. ~5.5cm gap between the plug and the food).

### Assays

#### Rapid iterative negative geotaxis (RING) assay

The RING assay set up was modified from an existing protocol (Gargano et al., 2005). The RING frame consisted of ~26 adjoining columns, ~1.2 cm wide and ~35 cm in height. The bottom of the frame was covered by doubled-over tape, to ensure a uniform base while ensuring that the surface is not sticky. This frame was loaded into a metallic support structure, consisting of two long rods to hold the frame in place, and a base covered by foam to absorb the shock, while maintaining it in a vertical position.

In each frame, 25 flies of one treatment and one sex were loaded into one column, and alternate columns were filled. 8 columns were assayed at a time in one round. Each such round had replicates from all treatment groups from one sex. Once the flies were loaded into the columns, the top was closed using cotton plugs, and the frame was mounted on the support. The flies were allowed to settle. The assay was performed in a dark room, with diffused light from the back of the set-up, to facilitate contrast for recording with a video camera (Sony HDR-PJ410).

The frame was mechanically disturbed, and moved sharply to the base, to make all the flies fall to the bottom. Once the flies were at the bottom, video recording was started, and a timer was kept for 30 seconds, which constituted one trial. After 30 seconds, the frame was disturbed similarly, and the process was repeated for 10 consecutive trials.

For each round, both the 1^st^ and the 10^th^ trial were scored. Screenshots were taken from the video recorded, at a fixed time point in the trials. The time-points were selected such that the snapshots were taken when the flies were dispersed throughout the set-up, and a majority of them had not reached the top. For males, this fixed point was 10s, while it was 15s for females.

The length of the column was divided into 31 bins of 1 cm. The number of flies in each bin were counted. If a fly was halfway between bins, it was counted in the bin in which its lowermost tip was present. The distance travelled was measured as the distance crossed by the entire body of the fly, that is, the lower limit of the bin in which it was scored.

Two parameters were scored – the average distance travelled by the flies of each treatment, and the propensity to show negative geotactic mobility. The propensity was measured as the total number of flies in each treatment that left the base of the set-up and travelled at least 1 cm. Being a fraction, the propensity data was arcsine-square root transformed before analysis (Zar, 1999).

#### Stop for sweet assay

##### Mechanical perturbation

The protocol was modified from Ries et al., (2017). After 3 days of stress (or control) treatment and recovery (as mentioned above), on the 4^th^ day, the flies were subjected to the stress (or control) protocol for 4 hours, but in the absence of water.

A cotton strip soaked in 99% glycerol was stuck across the middle of a 35mm petri plate of thickness 1.5cm. The plate was covered by a lid and sealed. Individual flies were aspirated into clean 5mm transparent glass tubes right before the assay. They were introduced into the set-up via a small hole drilled into the side of the lid, with the help of a glass tube and an aspirator. The fly was then shaken down to the bottom of the plate and allowed to wander around in the setup. For each fly, it was scored whether during a cross-over of the strip, it overran the glycerol or stopped to eat. Care was taken to only count the stops where the fly was eating, and not grooming. After each time the fly ran over the glycerol or stopped to eat, the setup was shaken again to let the fly start from the bottom of the plate. This process was repeated for 10 cross-overs for each fly.

Each set-up was scored at the time of the assay, by observers trained to identify the behaviours of stopping and feeding versus not-stopping, but blind to the nature of the treatment. The proportion of stops by each fly was calculated, given by:

(Number of times each fly stopped to eat) / (Total number of cross-overs monitored)

##### Adult crowding

After 72 hours of crowded (or control) conditions and 14 hours of recovery, both the treatment and control groups were subjected to 4 hours of starvation and desiccation. The assay was performed similarly as described above.

#### Exploratory behaviour assay

To measure exploratory tendency in flies, a previously reported experimental protocol (Soibam et al., 2012) was modified, and the activity was recorded using a video camera (Sony HDR-PJ410, Sony DCR-SR20E) for scoring later. The experimental arena consisted of a clear polycarbonate petri dish lid, with an inner diameter of 10 cm. The lid was placed over a blank sheet of paper having two concentric circles. The outer circle was of the same diameter as the lid, while the inner circle was such that it divided the arena into two zones – the zone between the outer and inner circle constituted 1/3rd of the total area, and the zone inside the inner circle constituted 2/3rd of the total area. Immediately before the assay, individual flies were aspirated into clean 5mm transparent glass tubes. They were introduced into the set-up with the help of the tube and an aspirator via a small hole drilled into the center of the lid. They were given 1 minute to acclimatize to their environment, and observed for the next 10 minutes.

As the flies tend to stay towards the outer edge of the arena, each time they entered the inner zone and came back was counted as one exploratory trip. The parameter scored was the total number of trips made by each fly within the 10-minute period.

#### Locomotor activity and rest

Locomotor activity of the flies was measured using *Drosophila* Activity Monitor (DAM2) data collection systems (Trikinetcs Inc., USA) using a standard protocol (Chiu et al., 2010). This system measures the activity of an individual fly in a glass tube as the number of times it crosses an infrared beam which bisects each channel in the DAM, perpendicular to the axis of the tube. Activity readings were taken every 5 minutes for a period of 6 hours.

After 3 days of stress and overnight recovery, flies were aspirated into transparent glass DAM tubes (5-mm diameter), devoid of any food, and plugged with cotton on each side. Aspiration was preferred over CO_2_ anaesthetization as the latter could affect their activity levels if the readings are taken without sufficient time for recovery from anaesthesia. The DAM tubes were loaded onto the monitors, with 32 flies in each monitor, and placed undisturbed in an incubator at 25°C at constant light.

The first 15 minutes of the data recorded was not scored to allow for acclimatization of the fly to the environment. Two parameters were scored for each fly – activity index and proportion of rest. Activity Index (AI) was measured as the total activity counts of a fly divided by the duration that the fly spent awake or not resting (Gilestro, 2012; Kayser et al., 2014). No activity for a period of 5 minutes was scored as rest (Chiu et al., 2010; Hendricks et al., 2000); the fraction of the assay duration spent resting was scored as the proportion of rest.

#### Starvation resistance assay

Following the recovery period after stress (or control) treatment, groups of 10 flies of each treatment and sex were made under light CO_2_ anaesthesia. They were transferred to vials containing 1.24% agar, which allowed for an environment of starvation but not desiccation. They were placed in an incubator at 25°C at constant light. At intervals of 4 hours following the set-up, the total number of flies alive in each vial were counted. This was continued till there were no flies alive in any vial.

Two parameters were scored – The Kaplan-Meier (KM) estimate (Kaplan and Meier, 1958) and the time point at which 50% of the flies in each vial died. The KM estimate for survival *S*(*t*) at time *t* was given by:

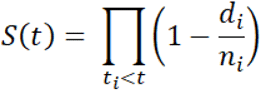

where *d_i_* is the number of flies that died at the time point *t_i_* and *n_i_* is the total number of flies which are at risk till just before the point *t_i_*.

### Statistical Analysis

Males and females were analysed separately for all the assays, because the stress treatment differed with sex.

For RING, replicates of treatment and control groups on which the assay was performed together were analysed together as one round. Two-factor mixed-model ANOVA was performed with treatment (stress or control) as a fixed factor, and round as a random factor. For all other assays, Mann-Whitney U (MWU) tests were performed with treatment (stress or control) as the factor, as the data failed Shapiro-Wilk normality tests. However, there were no major changes in significance levels of data when MWU test results were compared to ANOVA results for the same datasets and all interpretations remain essentially unchanged, which demonstrates the robustness of our results. Therefore, here we report only the results of the non-parametric MWU tests.

For all experiments, Cohen’s *d* effect sizes were estimated to compare between groups. The value of effect size was interpreted as large (*d* > 0.8), medium (0.8 > *d* > 0.5) or small (*d* < 0.5) following standard recommendations (Cohen, 1988). MWU tests were performed using Past3 (Hammer et al., 2001) and ANOVAs were performed using STATISTICA ver. 5 (StatSoft Inc). Survivorship curves for starvation resistance were plotted in SigmaPlot 11.0 (Systat Software Inc.) All other graphs were plotted in R version 3.1.3 (R Core Team, 2015).

## RESULTS

For all the experiments, the statistical data has been reported in Table 1.

**Table 1.**
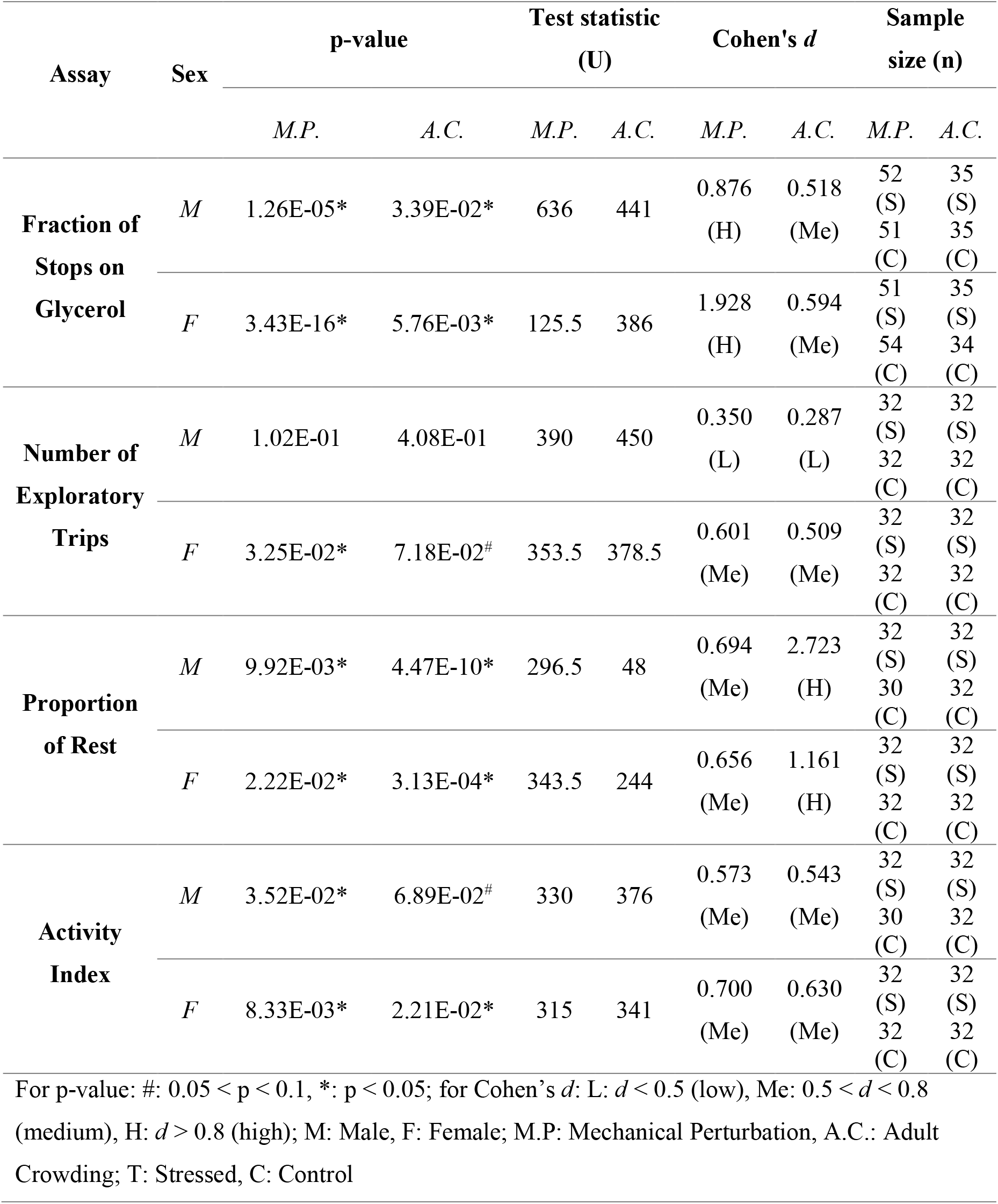
p-values, test statistics, Cohen’s *d* and sample sizes for various assays comparing behavioural responses in stressed versus control flies

| Assay | Sex | p-value |  | Test statistic (U) |  | Cohen's *d* |  | Sample size (n) |  |
| --- | --- | --- | --- | --- | --- | --- | --- | --- | --- |
|  |  | *M.P.* | *A.C.* | *M.P.* | *A.C.* | *M.P.* | *A.C.* | *M.P.* | *A.C.* |
| Fraction of Stops on Glycerol | *M* | 1.26E-05* | 3.39E-02* | 636 | 441 | 0.876 (H) | 0.518 (Me) | 52 (S) | 35 (S) |
|  |  |  |  |  |  |  |  | 51 (C) | 35 (C) |
|  | *F* | 3.43E-16* | 5.76E-03* | 125.5 | 386 | 1.928 (H) | 0.594 (Me) | 51 (S) | 35 (S) |
|  |  |  |  |  |  |  |  | 54 (C) | 34 (C) |
| Number of Exploratory Trips | *M* | 1.02E-01 | 4.08E-01 | 390 | 450 | 0.350 (L) | 0.287 (L) | 32 (S) | 32 (S) |
|  |  |  |  |  |  |  |  | 32 (C) | 32 (C) |
|  | *F* | 3.25E-02* | 7.18E-02<sup>#</sup> | 353.5 | 378.5 | 0.601 (Me) | 0.509 (Me) | 32 (S) | 32 (S) |
|  |  |  |  |  |  |  |  | 32 (C) | 32 (C) |
| Proportion of Rest | *M* | 9.92E-03* | 4.47E-10* | 296.5 | 48 | 0.694 (Me) | 2.723 (H) | 32 (S) | 32 (S) |
|  |  |  |  |  |  |  |  | 30 (C) | 32 (C) |
|  | *F* | 2.22E-02* | 3.13E-04* | 343.5 | 244 | 0.656 (Me) | 1.161 (H) | 32 (S) | 32 (S) |
|  |  |  |  |  |  |  |  | 32 (C) | 32 (C) |
| Activity Index | *M* | 3.52E-02* | 6.89E-02<sup>#</sup> | 330 | 376 | 0.573 (Me) | 0.543 (Me) | 32 (S) | 32 (S) |
|  |  |  |  |  |  |  |  | 30 (C) | 32 (C) |
|  | *F* | 8.33E-03* | 2.21E-02* | 315 | 341 | 0.700 (Me) | 0.630 (Me) | 32 (S) | 32 (S) |
|  |  |  |  |  |  |  |  | 32 (C) | 32 (C) |

For p-value: #: $0.05 < p < 0.1$ , \*: $p < 0.05$ ; for Cohen's *d*: L: $d < 0.5$ (low), Me: $0.5 < d < 0.8$ (medium), H: $d > 0.8$ (high); M: Male, F: Female; M.P: Mechanical Perturbation, A.C.: Adult Crowding; T: Stressed, C: Control

### No Change in Innate Response

There were no significant differences between the stressed flies and the controls of either sex in either propensity of negative geotaxis, or their ability to climb the walls of the RING setup (Figs S1–S4). This indicates that neither stressor injured or caused physical harm to the flies, as negative geotaxis, a cue-based response, is unchanged (Gargano et al., 2005).

### Lesser Interest in Pleasurable Activities

Flies subjected to mechanical perturbation showed significantly reduced tendency to feed on glycerol as compared to their controls across both males (Fig. 1A) and females (Fig. 1B). This indicates that this stressor induced anhedonia, i.e. a reduction of interest in pleasurable activities.

**Fig. 1.**
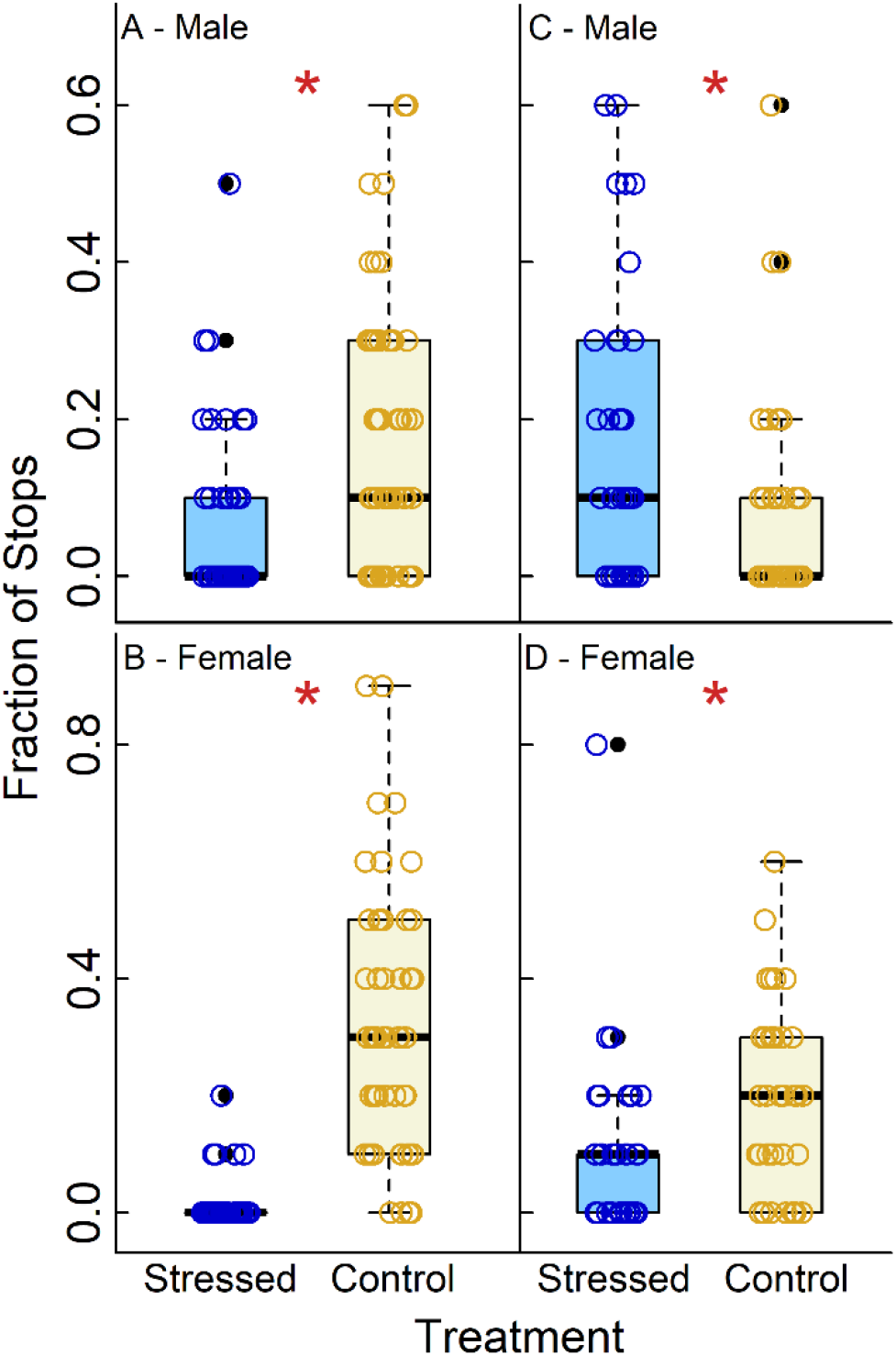
Anhedonic response to stress. Fraction of stops to feed on glycerol in A. males and B. females after mechanical perturbation; C. males and D. females after adult crowding vs their respective controls. * indicates MWU p < 0.05. The points represent the data for all replicates of the particular group with small random jitter on the x-axis. The edges of the box denote the 25^th^ and 75^th^ percentiles, the black solid line represents the median. The whiskers extend to the extreme data point, which is no more than 1.5 times the inter-quartile range from the top or bottom of the box. The points beyond this are indicated as outliers (solid black circles).

When subjected to adult crowding, female flies showed anhedonia and fed lesser on glycerol (Fig. 1D). However, male flies showed an increased tendency to feed on glycerol (Fig. 1C). This suggests that with respect to anhedonic behaviour, crowding induces sexual dimorphism in flies.

### Reduced Exploration of Novel Habitat in Females

Male flies showed no significant change in the tendency to explore their habitat after mechanical perturbation (Fig. 2A). However, there was a significant reduction in the number of exploratory trips made by the female flies (Fig. 2B) subjected to this stressor. Also, although adult crowding did not lead to significant change in exploratory tendency of the males (Fig. 2C), there was an almost significant (p < 0.1) reduction in the number of exploratory trips in females, with a medium effect size (Fig. 2D, Table 1).

**Fig. 2.**
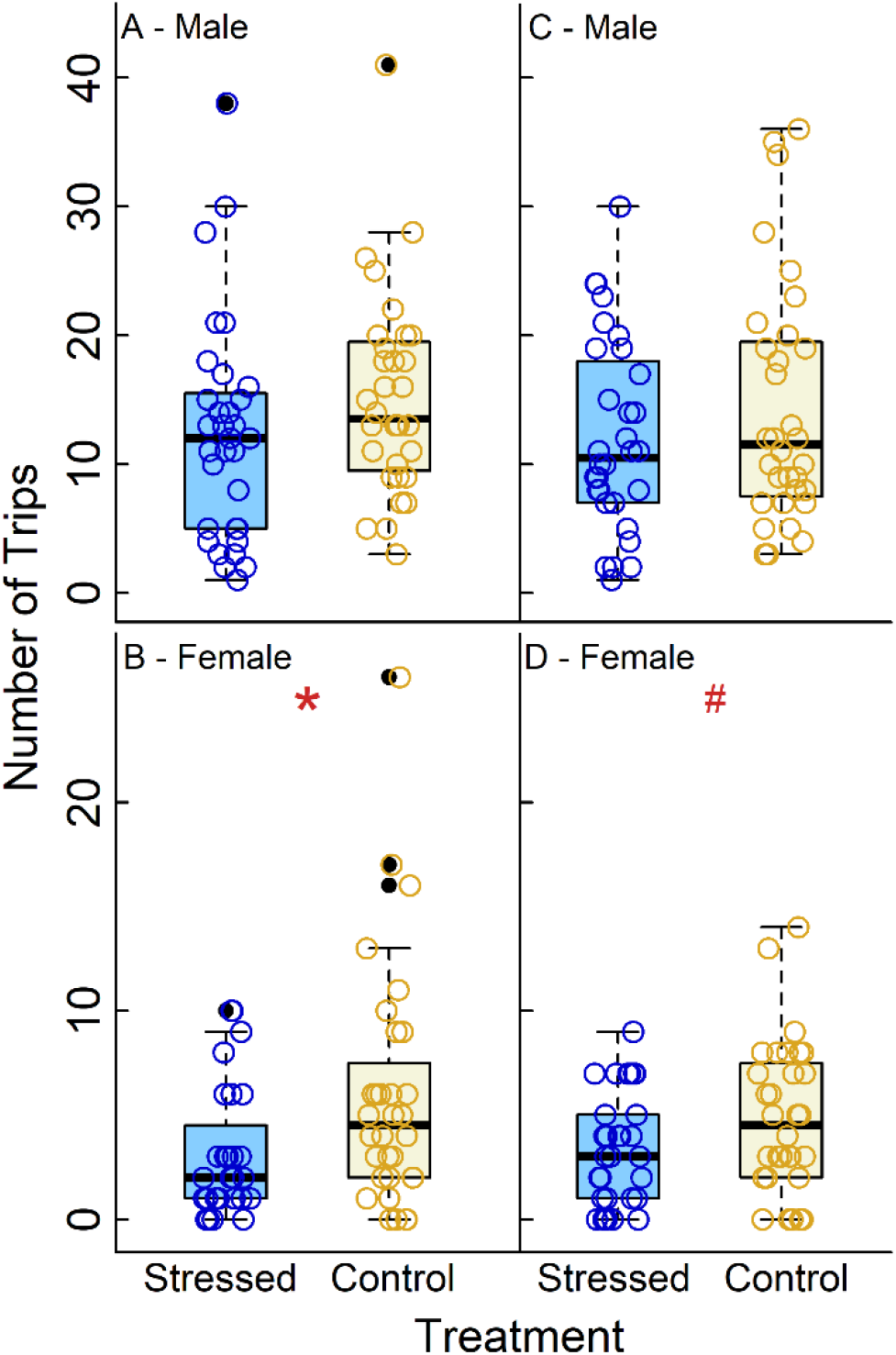
Changes in exploratory tendency due to stress: Number of exploratory trips in A. males and B. females after mechanical perturbation; C. males and D. females after adult crowding vs their respective controls. * indicates MWU p < 0.05; # indicates p < 0.1.

Taken together, it can be stated that our stressors do not affect the exploratory tendencies of male flies, but reduces the same for the female flies. This suggests that stress induces a sexual dimorphism in exploratory behaviour in fruit flies.

### Increased Locomotor Activity and Insomnia

The proportion of time spent resting was significantly lowered after mechanical perturbation in both males (Fig. 3A) and females (Fig. 3B). Similar reduction was also observed across both sexes after adult crowding (Figs 3C & 3D).

**Fig. 3.**
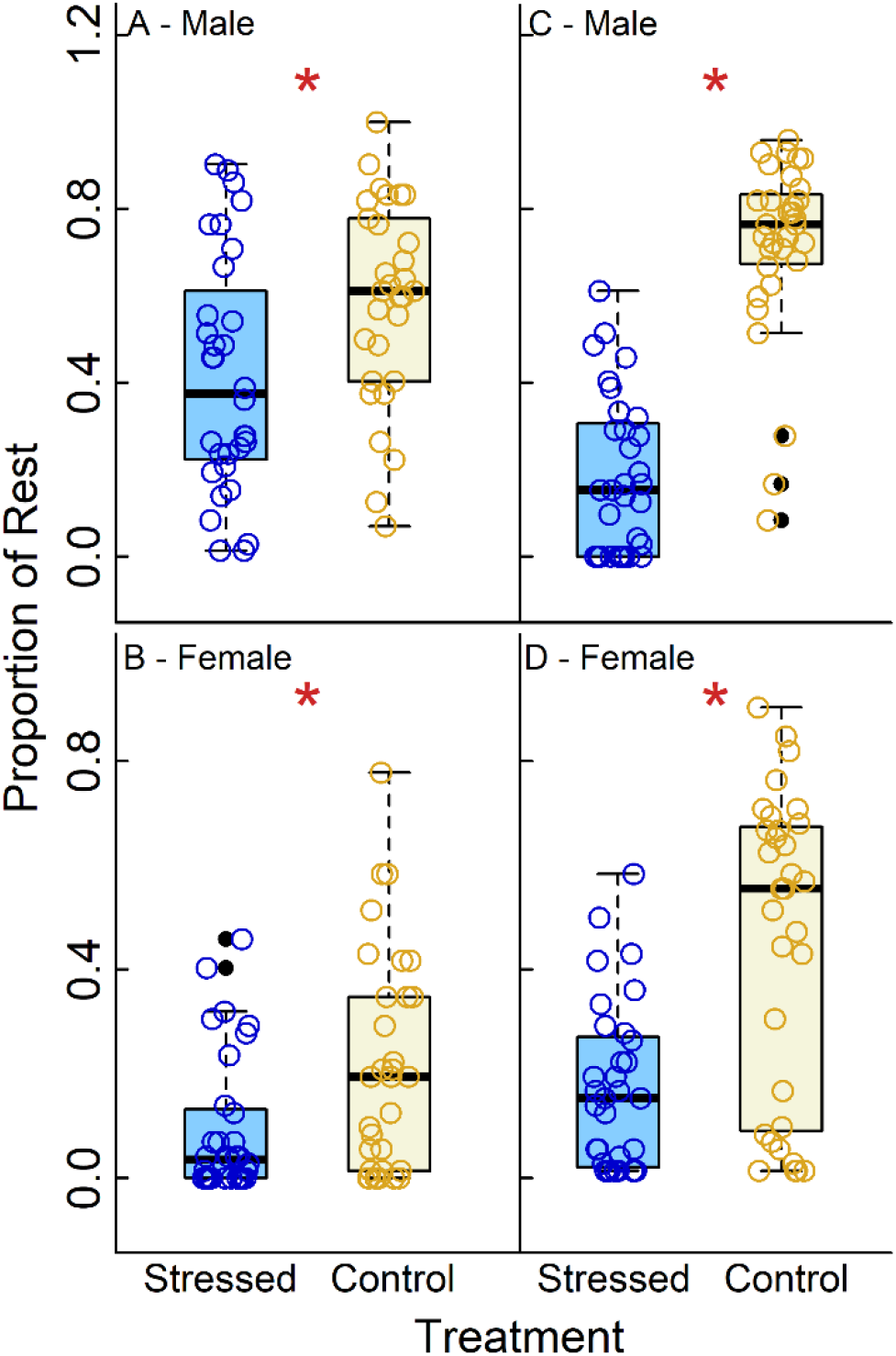
Stress-induced changes in sleep or rest levels: Proportion of time spent resting over 6 hours in A. males and B. females after mechanical perturbation; C. males and D. females after adult crowding vs their respective controls. * indicates MWU p < 0.05.

However, when the Activity Index (AI) was compared for these stressors, crowding again induced a sexual dimorphism. While both males (Fig. 4A) and females (Fig. 4B) showed increased AI after mechanical perturbation, after crowding, males showed an almost significant increase in AI (Fig. 4C), while females showed a reduction in AI (Fig. 4D).

**Fig. 4.**
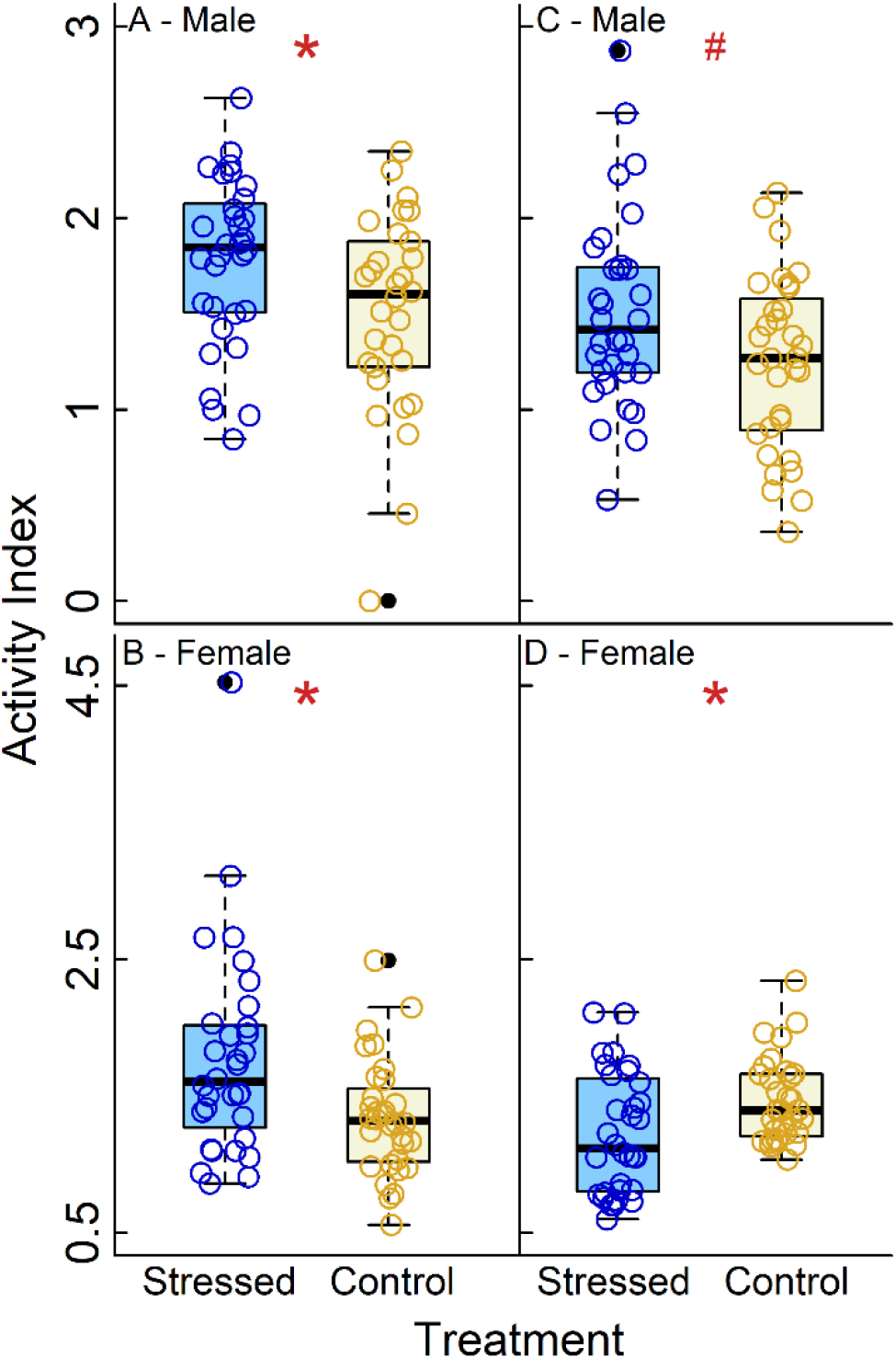
Effect of stress on activity during wakefulness: Activity Index over 6 hours in A. males and B. females after mechanical perturbation; C. males and D. females after adult crowding vs their respective controls. * indicates MWU p < 0.05, # indicates p < 0.1

Thus, while stress makes flies rest less across sexes, the nature of stressor modulates sexual dimorphism in AI levels.

### No Change in Starvation Resistance

When the starvation resistance of flies which had been subjected to adult crowding was compared to their controls, there was no difference in the time taken for 50% mortality in the vial across treatment and control groups for both males and females (Table S1). This is congruent with the observation that the KM survivorship curves almost superimpose in both cases (Fig. S5).

Fig. 5 presents a schematic summary of all the results.

**Fig. 5:**
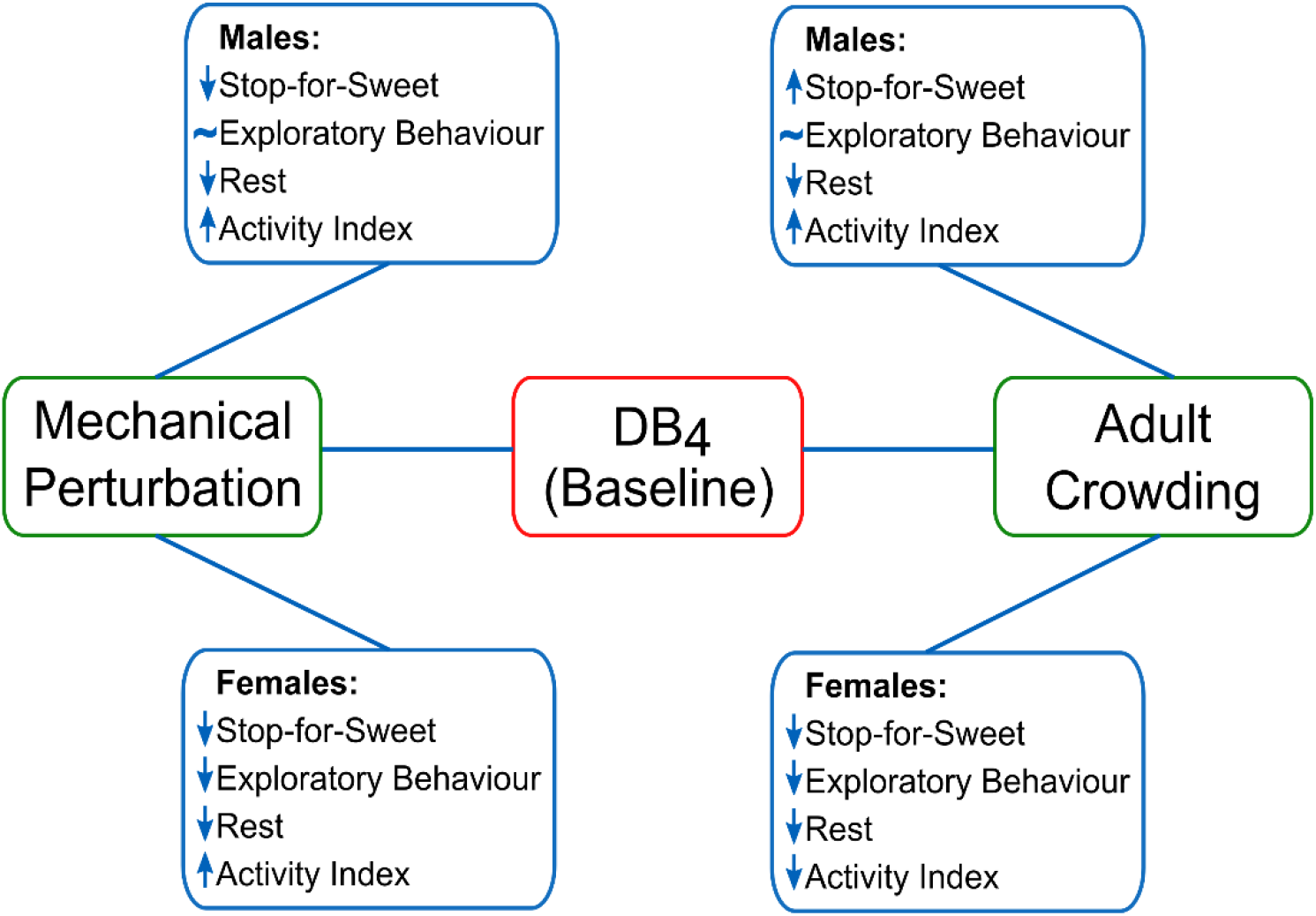
Summary of the experimental results. Behavioural changes due to different stressors in male and female baseline flies. ↑ indicates increase in level of the behaviour measured in stressed flies vs. their controls, ↓ indicates decrease in the level of the behaviour measured in stressed flies vs. their controls, ~ indicates no difference in the level of the behaviour measured between the stressed and control flies.

## DISCUSSION

### Nature of Stressor Modulates Sexual Dimorphism in Anhedonic Behaviour

Hedonic behaviours have been widely used to measure stress-response in rodent models of Chronic Mild Stress (CMS) (see Willner, 2017; Willner et al., 1992 for discussions). In this paradigm, the animals are subjected to a series of unpredictable, mild, largely abiotic stressors over several weeks (Willner, 2017). This typically leads to a reduced preference to feed on sucrose which is considered anhedonic, i.e. indicating a lack of interest in a pleasurable activity (Katz, 1982; Willner et al., 1987). Similarly, in male *D. melanogaster*, a variety of abiotic stresses have been shown to induce anhedonia, which has been measured as a reduction in feeding on glycerol (Ries et al., 2017) or sucrose (Araujo et al., 2018).

In our experiments, the abiotic mechanical perturbation stress paradigm led to a reduction in glycerol feeding in both male and female flies, thus indicating a lack of motivation to partake in pleasurable activities (Figs 1A & 1B). However, the biotic stressor – adult crowding – affected male and female flies differently. This result is consistent with previous studies in rodents, where social instability, with periods of isolation and crowding, resulted in sex-specific anhedonic responses (Herzog et al., 2009).

In our experiments, adult crowding induced anhedonia only in stressed females (Fig. 1D), while counter-intuitively, stressed males showed an increase in glycerol feeding (Fig. 1C). A possible reason for this could have been that crowding was leading to a competition for resources (Joshi and Mueller, 1997). Since male flies have lower body size, they could have been affected more severely by reduced access to food under crowded condition, thus leading effectively to starvation. This starvation could then be providing an immediate impetus for the male flies to feed during the stop-for-sweet assay. On the other hand, female flies are larger than the males and are known to be more resistant to starvation (Chippindale et al., 1996; Zwaan et al., 1991). Thus, the females could be responding primarily to the biotic stressor in a similar way as the abiotic one, thus exhibiting a reduced motivation state, i.e. anhedonia. Stated differently, it was possible that the males were hungrier (which trumped anhedonia) while the females were less hungry and therefore exhibited anhedonic symptoms. To investigate this possibility, we assayed the starvation resistance of the stressed and unstressed flies. We found that adult crowding does not have an effect on the starvation resistance of either males or females (Fig. S5), thus overruling this possibility. Thus, the physiological reason for this dimorphism remains unclear. Summarily, it can be stated that the nature of the stressor plays a crucial role in anhedonic responses to stress, and sexual dimorphism in sex response seems to be modulated by the nature of the stressor. To the best of our knowledge, this is the first demonstration of a sexually dimorphic anhedonic response to stress in *D. melanogaster*. Anhedonia is a key diagnostic symptom of depression in human beings, and is known to be sexually dimorphic in humans (Fonseca-Pedrero et al., 2008) and rodents (Lu et al., 2015).

### Stress Reduces Motivation to Explore Novel Habitat in Females

Our paradigm of non-lethal 3-day stressors revealed a sexual dimorphism in exploratory behaviour in response to stress. Male flies showed no change in the number of exploratory trips, while female flies explored significantly lesser. This dimorphism was consistent across both the biotic and abiotic stressor (Fig. 2). This is in keeping with previous results of sexual dimorphism in this behaviour in flies after 24-hour long starvation and oxidative stress (Neckameyer and Matsuo, 2008).

The tendency to explore is related to seeking out novel habitats (Cote et al., 2010) and is also energy intensive. This decrease in exploratory tendencies of female flies after stress could indicate either a physical inability to explore due to exhaustion or injury from the stressor, or a lack of motivation to explore new surroundings, or both. However, it is crucial to note that the cue-based response of negative geotaxis is not affected across sexes by either stressor (Figs S1–S4), indicating that the changes are not likely due to physical harm, fatigue or injury to the fly. Thus, we conclude that these flies lack the motivation to explore after being stressed. Additionally, preference for edges in flies is postulated to be a marker of seeking shelter (Liu et al., 2007), and the increase in this behaviour could possibly represent increased fear or anxiety-like behaviour due to stress.

Exploratory behaviour is related to locomotor activity levels in rats (Willig et al., 1987). In fruit flies, exploration is characterized by an initial elevated level of activity (Liu et al., 2007). Hence, we next investigated the impact of stress on locomotor activity.

### Stress Causes Insomnia Across Sexes and Changes Locomotor Activity

Long-term changes in rest and activity patterns is indicative of a lasting effect of stress on the organism. After sufficient time to acclimatize to the environment of recording (see Materials and Methods), over the next 6-hours, we observed that stress-induced a change in locomotor activity and rest levels. Both mechanical perturbation and crowding caused the flies to spend lesser time resting or sleeping across sexes (Fig. 3). Our sleep results in flies are congruent with previous observations in rats that suggest disruption of sleep patterns after CMS (Cheeta et al., 1997)

Activity Index (AI) is a measure of the flies’ activity in the DAM tube during their period of wakefulness (Gilestro, 2012; Kayser et al., 2014). Mechanical perturbation resulted in increased locomotor activity during wakefulness in both males and females (Fig. 4A & 4B). This observation, coupled with lower rest levels, indicates that this stressor induces hyperactivity in flies. However, adult crowding brings about sexual dimorphism in activity indices of flies. While male flies subjected to this stressor showed hyperactivity (Fig 4C), crowded female flies were less active than their controls when awake (Fig. 4D). The increased activity in male flies over 6 hours is in contrast to previous studies in flies subjected to vibrational stress, which found a reduction in locomotor activity in stressed males over a 15-minute period (Ries et al., 2017). Thus, the differences in the observations could be due to the vastly different durations over which activity has been measured in the two studies. Our recordings, over a considerably longer duration, allow for an initial acclimatization period to the environment. Thus, the immediate exploratory activity in a new environment is excluded from the data, allowing for basal changes in psychomotor activity and rest levels to be studied.

While exploration and locomotor activity seem to be correlated (Liu et al., 2007; Willig et al., 1987), our results suggest that stress can impact these two behaviours in very different ways. Higher activity levels and reduced rest over 6 hours in males after stress does not cause a concomitant increase in exploratory activity. Rather, a decrease in exploratory behaviour in females occurs, which can be interpreted as a measure of a reduced motivational state.

## IMPLICATIONS

In this study, we showed that stressors of differing nature (biotic vs abiotic) can cause varying behavioural responses, and these can be modulated by the sex of the fly. These results have several interesting implications. Our results are congruent with observations in human beings that males and females differ in terms of their propensity of various mood disorders (Kessler, 2003; McLean et al., 2011). For example, in human beings, it is known that males show larger predisposition to alcoholism and other drug abuse, antisocial personality disorder and attention deficit disorders while depression, anxiety and eating disorders predominate in females (see Palanza, 2001 for a discussion). Further, social contexts for male and female animals are innately different due to differences in their social roles, differential parental investment etc. Thus, social stresses are more likely to induce sexually dimorphic responses (see Palanza and Parmigiani, 2017 for a discussion). This notion is supported by our observation that adult crowding, which is a biotic stressor, induces sexual dimorphism in a larger number of behaviours. We also found that in general, female flies were more affected by the stressors than the male flies. This is consistent with human data on SIMDs that suggest that women are more prone to depression (Kessler, 2003; Nolen-Hoeksema, 1987) and anxiety (McLean et al., 2011; Wittchen et al., 1994).

These results thus strengthen the case for using *D. melanogaster* as a model system to investigate the phenomenon of sex differences in SIMDs. Previous studies have suggested that sexual dimorphism in response to acute stress in *D. melanogaster* is modulated by sex-specific hormones, and the hormonal environment of the brain can determine which neurons are recruited into the stress-response circuitry (Neckameyer and Matsuo, 2008). This could potentially cause the observed sexually dimorphic responses to chronic stress as well. More critically, if one can show a reasonable degree of convergence between humans and fruit flies in terms of the genetic and physiological mechanisms underlying these disorders, then a lot of research on sex differences in SIMDs can shift to the *Drosophila* system. The advantages of this model system, in terms of genetic, neurobiological and behavioural tractability, can allow for a detailed understanding of this dimorphism. Given that *Drosophila* has already proven to be a good system to model Alzheimer’s disease, cardiovascular disease and diabetes, to name a few (reviewed in Pandey and Nichols, 2011), the possibility of furthering research on sexual dimorphism in SIMDs using the fruit fly is rather tantalizing.

## LIST OF SYMBOLS AND ABBREVIATIONS

SIMD: Stress-Induced Mood Disorder
RING: Rapid Iterative Negative Geotaxis
DAM: *Drosophila* Activity Monitor
AI: Activity Index
KM: Kaplan-Meier
MWU: Mann-Whitney U
CMS: Chronic Mild Stress

## ACKNOWLEDGMENTS

We thank Nikita Sabnis, Radha Barve and Sandesh Papade for help in running the experiments, and Sudipta Tung and Abhishek Mishra for insightful comments. This work was supported by a research grant (#EMR/2014/000476) from Science and Engineering Research Board (SERB), Department of Science and Technology, Government of India and internal funding from IISER-Pune.

## AUTHOR CONTRIBUTIONS

S.L. and S.D. formulated the study, S.L., A.M._1_, S.S., A.M._2_, A.T. and G.K. carried out the experiments, S.L., A.M._1_ and S.S. analysed the data, S.L. and S.D. wrote the manuscript with inputs from other authors.

## DATA AVAILABILITY

Data will be deposited in Dryad if the manuscript is accepted for publication.

## SUPPLEMENTARY FIGURES AND INFORMATION

### No Changes in Innate Responses Due to Stress

Compared to their controls, neither male (Fig. S1A) nor female (Fig. S1B) flies subjected to mechanical perturbation showed any significant change in their propensity of negative geotaxis measured in the 1^st^ trial of the RING assay. Similar results were obtained for males (Fig. S1C) and females (Fig. S1D) subjected to adult crowding.

Similarly, neither males (Fig. S2A) nor females (Fig. S2B) showed a change in the average distance travelled during negative geotaxis after mechanical perturbation. These trends were also retained when male (Fig. S2C) and female (Fig. S2D) flies were subjected to adult crowding.

When measured after 10 trails, no change in propensity of negative geotaxis was observed across both sexes after mechanical perturbation (Figs S3A & S3B) or after adult crowding (Figs S3C & S3D).

No changes were observed in the average distance travelled by males or females after mechanical perturbation (Figs S4A & S4B) and after crowding (Figs S4C & S4D).

**Fig. S1.**
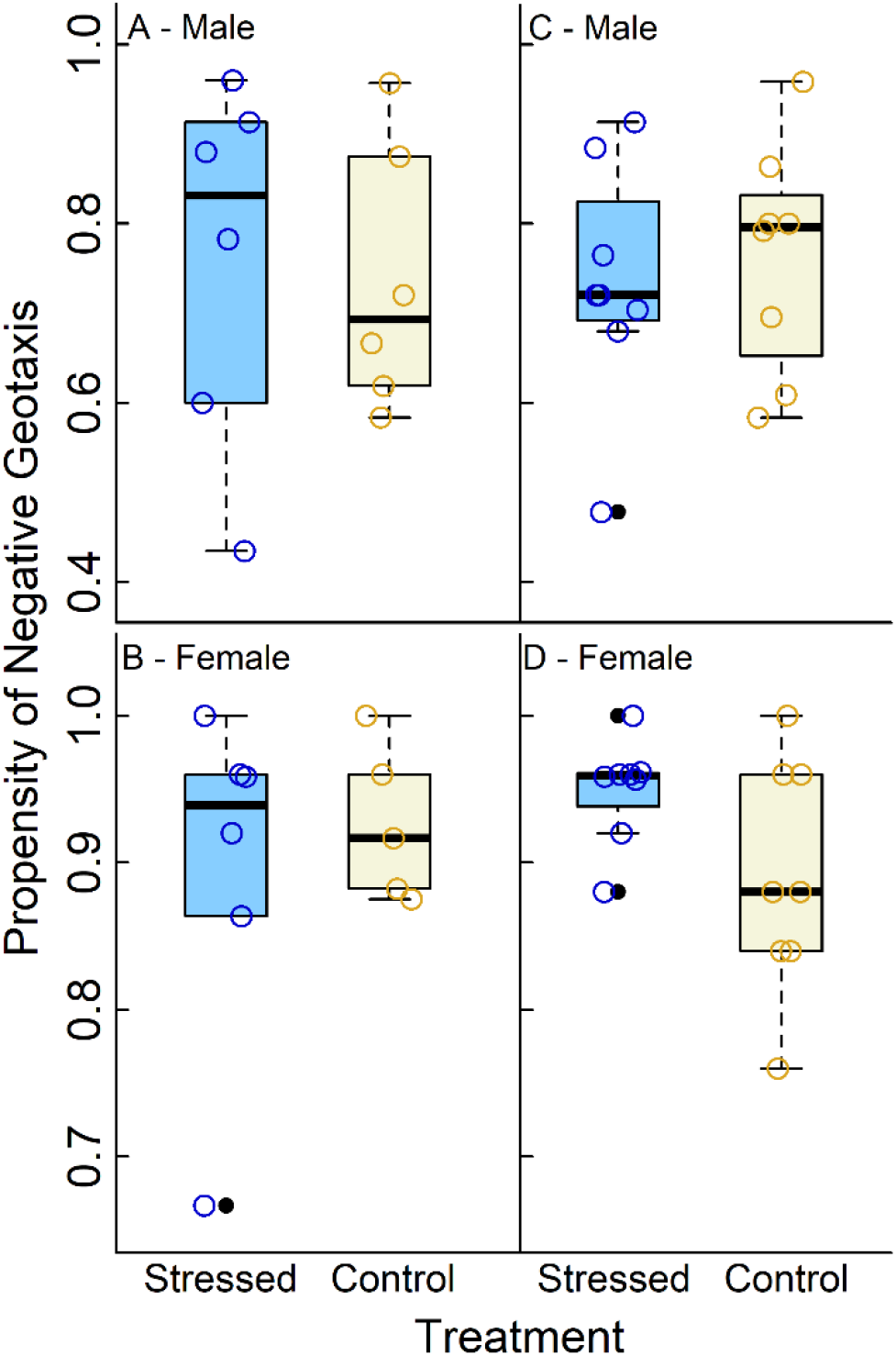
No effect of stress on negative geotactic propensity: Propensity of negative geotaxis after the 1st trial in A. males and B. females after mechanical perturbation; C. males and D. females after adult crowding vs their respective controls

**Fig. S2:**
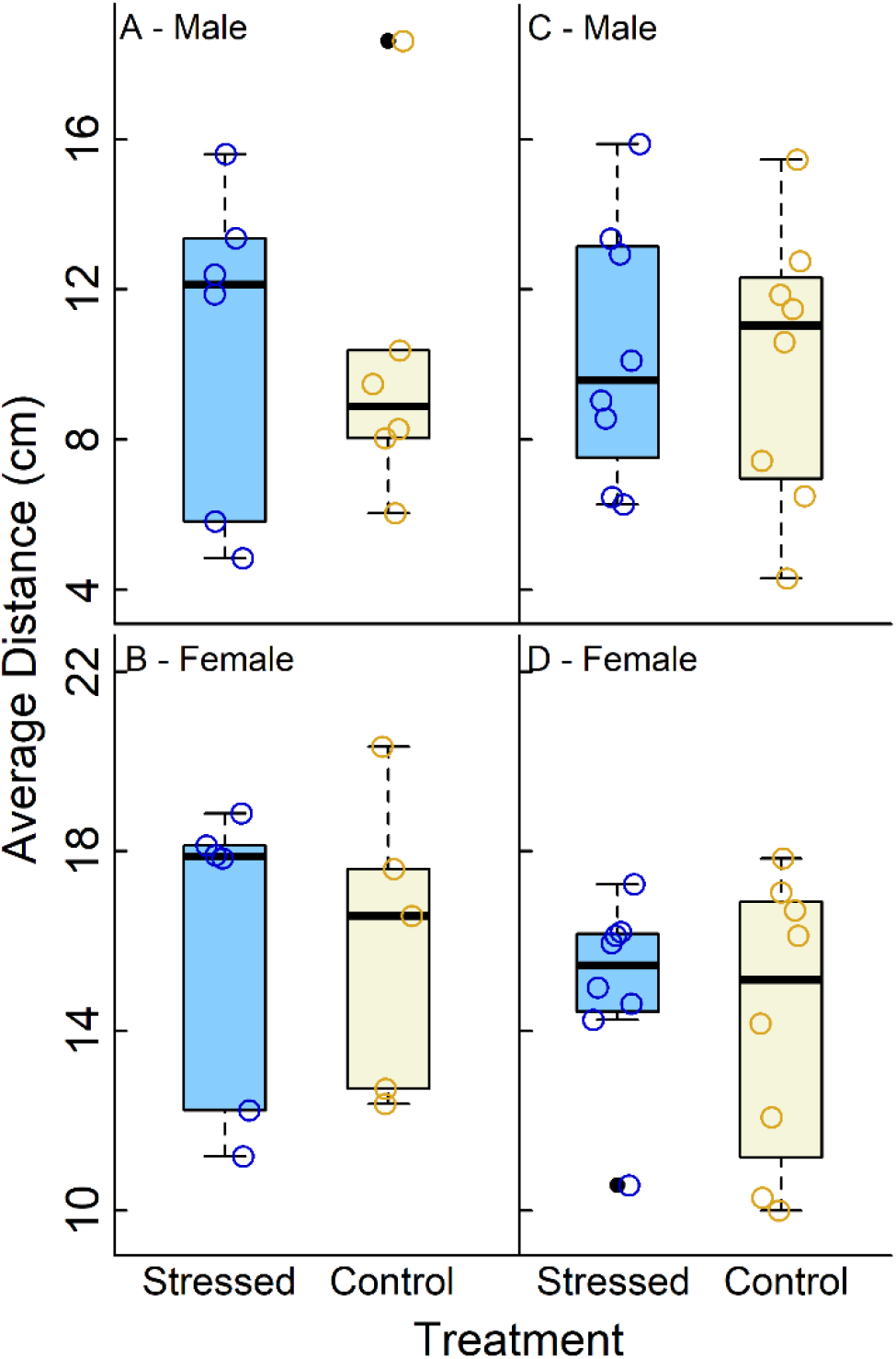
No effect of stress on negative geotactic distance travelled: Average distance travelled after the 1st trial in A. males and B. females after mechanical perturbation; C. males and D. females after adult crowding vs their respective controls

**Fig. S3.**
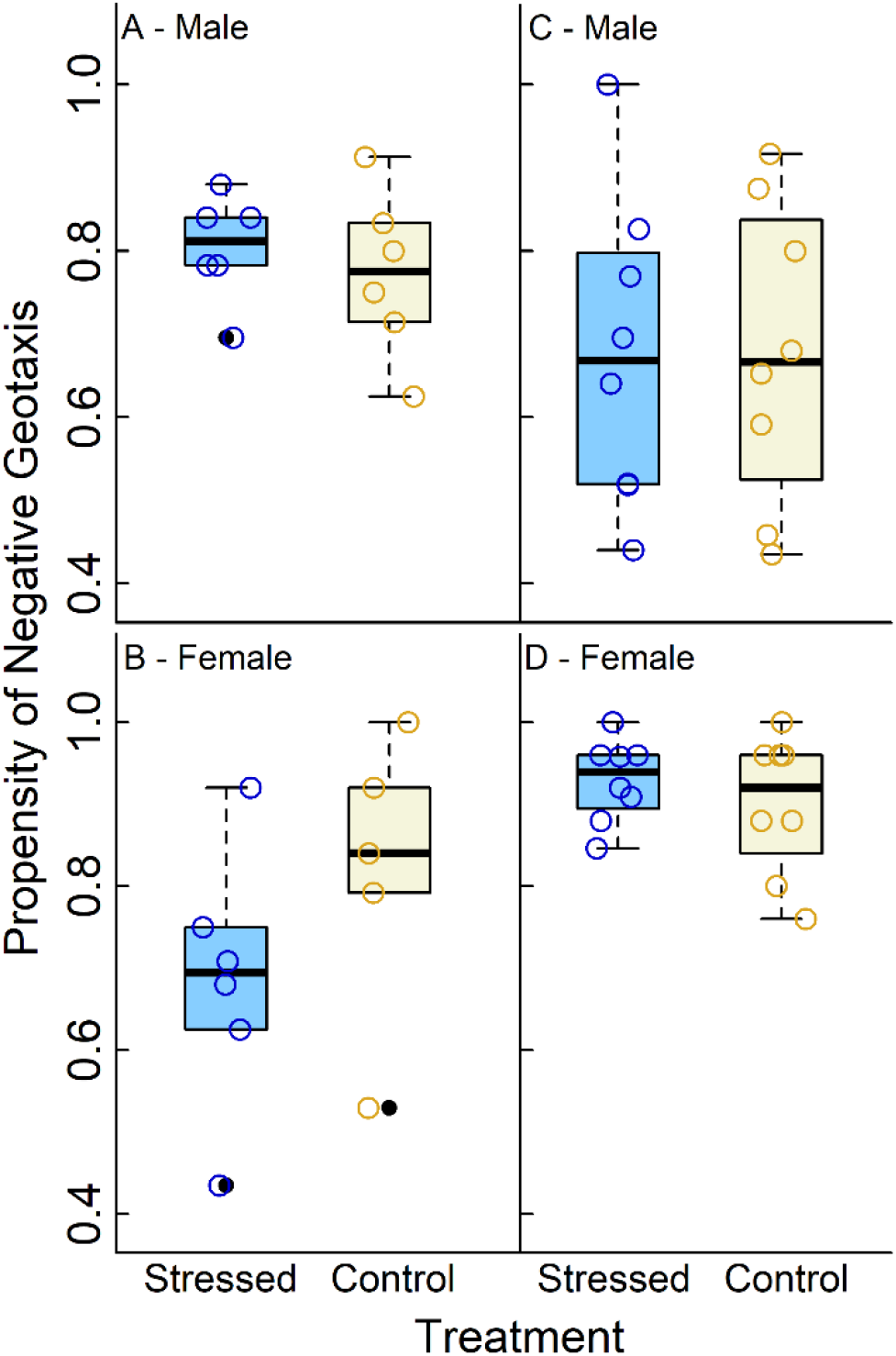
No changes in negative geotactic propensity after several trials: Propensity of negative geotaxis after 10 trials in A. males and B. females after mechanical perturbation; C. males and D. females after adult crowding vs their respective controls

**Fig. S4.**
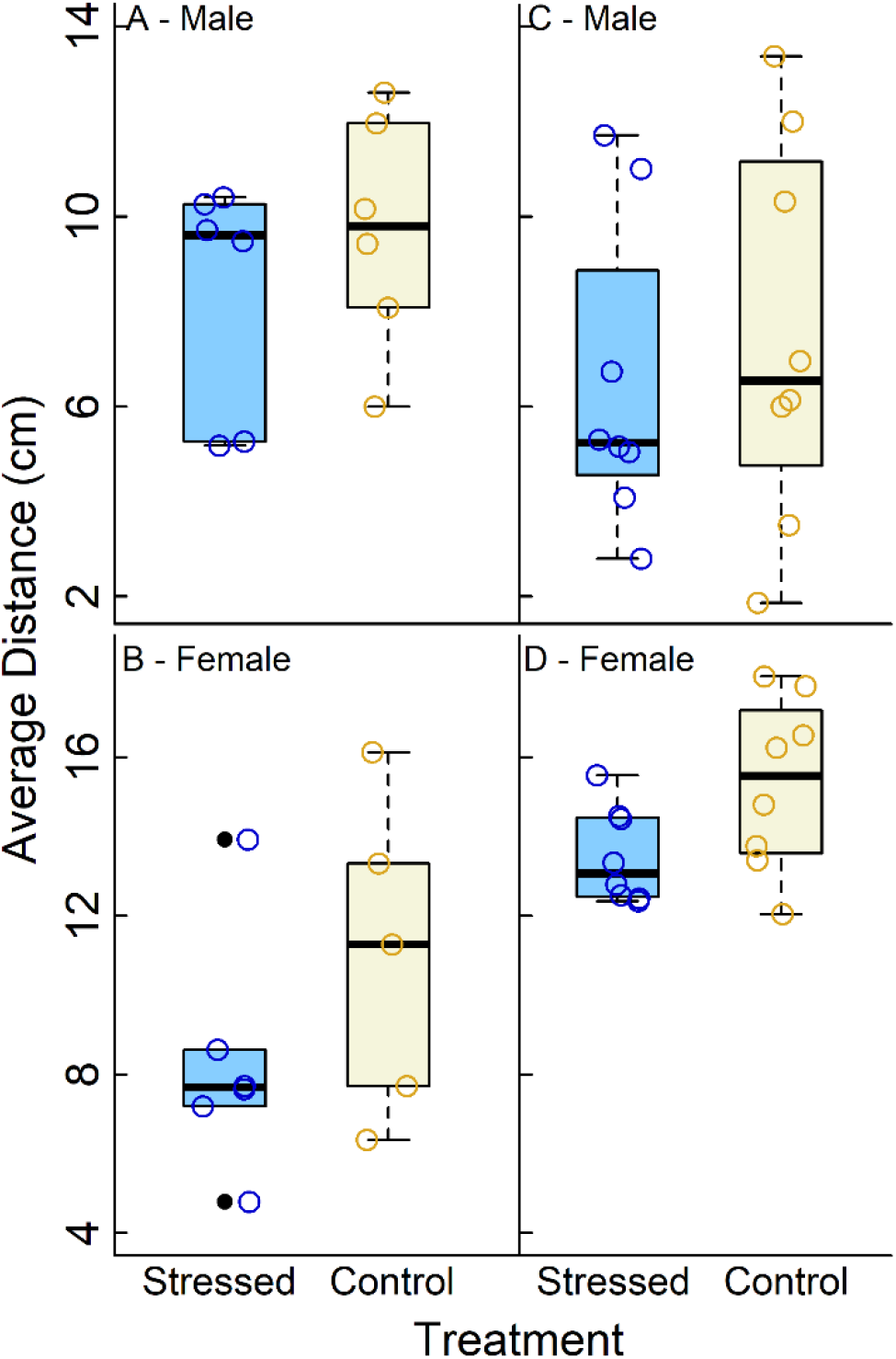
No changes in negative geotactic distance after several trials: Average distance travelled after 10 trials in A. males and B. females after mechanical perturbation; C. males and D. females after adult crowding vs their respective controls

**Fig. S5.**
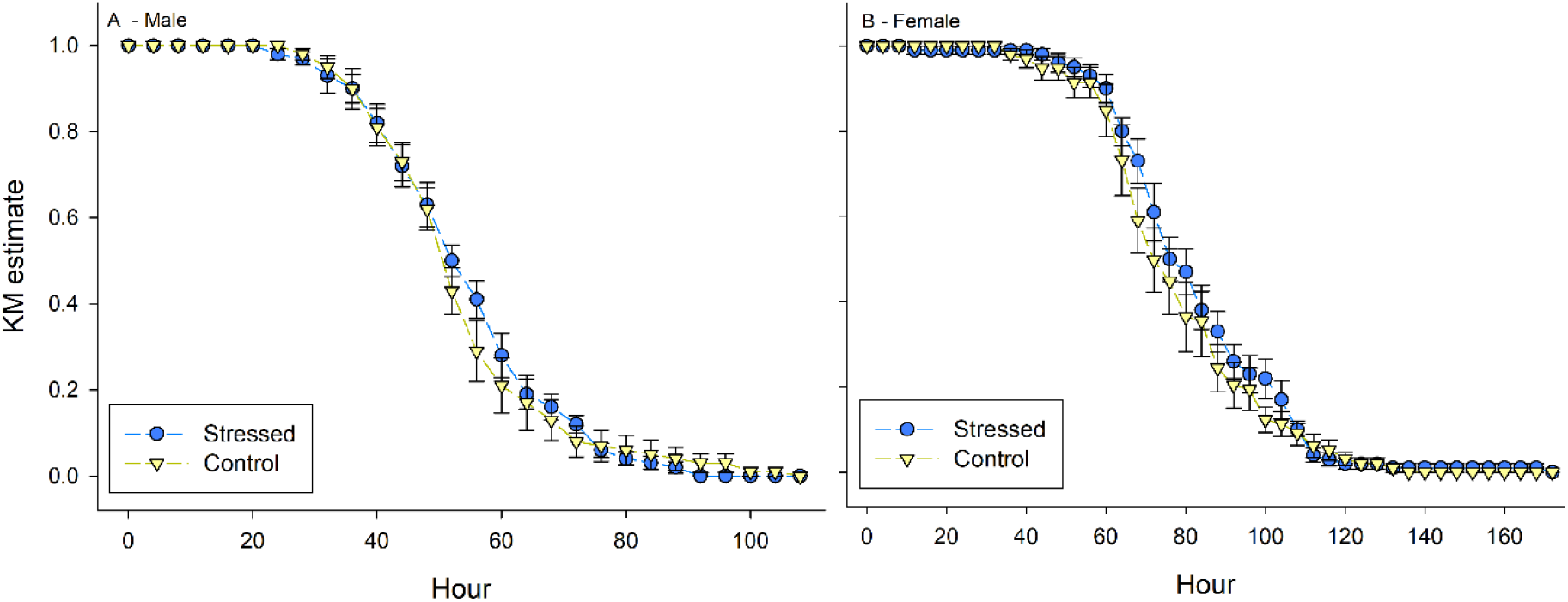
No change in starvation resistance after adult crowding: Survivorship curve under starvation conditions based on KM estimates for A. males and B. females after adult crowding compared to their respective controls.

**Table S1.** p-values, test statistics, Cohen’s *d* and sample sizes for the RING and starvation resistance assays comparing stressed versus control flies

| Assay | Sex | p-value |  | Test statistic |  | Cohen's *d* |  | Sample size (n) |  |
| --- | --- | --- | --- | --- | --- | --- | --- | --- | --- |
|  |  | M.P. | A.C. | M.P. | A.C. | M.P. | A.C. | M.P. | A.C. |
| **RING Propensity (Trial 1)** | M | 0.512 | 0.663 | F(1,2) = 0.62 | F(1,2) = 0.23 | 0.140<br>(L) | 0.227<br>(L) | 6<br>(S) | 8<br>(S) |
|  |  |  |  |  |  |  |  | 6<br>(C) | 8<br>(C) |
|  | F | 0.670 | 0.335 | F(1,2) = 0.24 | F(1,2) = 2.96 | 0.343<br>(L) | 0.970<br>(H) | 6<br>(S) | 8<br>(S) |
|  |  |  |  |  |  |  |  | 5<br>(C) | 8<br>(C) |
| **RING Average Distance (Trial 1)** | M | 0.637 | 0.893 | F(1,2) = 0.30 | F(1,2) = 0.02 | 0.117<br>(L) | 0.079<br>(L) | 6<br>(S) | 8<br>(S) |
|  |  |  |  |  |  |  |  | 6<br>(C) | 8<br>(C) |
|  | F | 0.924 | 0.386 | F(1,2) = 0.01 | F(1,2) = 2.08 | 0.033<br>(L) | 0.268<br>(L) | 6<br>(S) | 8<br>(S) |
|  |  |  |  |  |  |  |  | 5<br>(C) | 8<br>(C) |
| **RING Propensity (Trial 10)** | M | 0.302 | 0.577 | F(1,2) = 1.90 | F(1,2) = 0.39 | 0.367<br>(L) | 0.001<br>(L) | 6<br>(S) | 8<br>(S) |
|  |  |  |  |  |  |  |  | 6<br>(C) | 8<br>(C) |
|  | F | 0.123 | 0.203 | F(1,2) = 6.68 | F(1,2) = 9.18 | 0.768<br>(Me) | 0.417<br>(L) | 6<br>(S) | 8<br>(S) |
|  |  |  |  |  |  |  |  | 5<br>(C) | 8<br>(C) |
| **RING Average Distance (Trial 10)** | M | 0.165 | 0.204 | F(1,2) = 4.61 | F(1,2) = 2.62 | 0.537<br>(Me) | 0.285<br>(L) | 6<br>(S) | 8<br>(S) |
|  |  |  |  |  |  |  |  | 6<br>(C) | 8<br>(C) |
|  | F | 0.194 | 0.192 | F(1,2) = 3.72 | F(1,2) = 10.34 | 0.745<br>(Me) | 1.045<br>(H) | 6<br>(S) | 8<br>(S) |
|  |  |  |  |  |  |  |  | 5<br>(C) | 8<br>(C) |
| **Starvation Resistance-50% mortality time** | M | ----- | 0.88 | ----- | U = 47.5 | ----- | 0.118<br>(L) | ----- | 10<br>(S) |
|  |  |  |  |  |  |  |  | ----- | 10<br>(C) |
|  | F | ----- | 0.59 | ----- | U = 42.5 | ----- | 0.265<br>(L) | ----- | 10<br>(S) |
|  |  |  |  |  |  |  |  | ----- | 10<br>(C) |

For p-value: #: $0.05 < p < 0.1$ , \*: $p < 0.05$ ; for Cohen's *d*: L: $d < 0.5$ (low), Me: $0.5 < d < 0.8$ (medium), H: $d > 0.8$ (high); M: Male, F: Female; M.P: Mechanical Perturbation, A.C.: Adult Crowding, S: Stressed; C: Control

